# A guanosine metabolism-bioenergetics intersection drives Parkinson’s disease

**DOI:** 10.64898/2026.07.29.741503

**Authors:** Alexandros C. Kokotos, Santiago R. Unda, Paula Reyes-Pérez, Mariam Isayan, Maria Teresa Periñan, Michael G. Kaplitt, Timothy A. Ryan

## Abstract

Several Parkinson’s Disease (PD) linked mutations are known to drive deficits in a pathway that relies on an adequate supply of guanosine nucleotide triphosphate (GTP) to drive cellular neuronal processes. Similarly, there is strong evidence that a deficit in bioenergetic support for neuron function is also a major genetic driver of PD. We show here that the reliance on these two purine-based metabolites intersect at another PD susceptibility gene that encodes nucleoside diphosphate kinase (NDK) which converts ATP into GTP. We show that overexpression of NDK is strongly protective both in-vivo and in-vitro to metabolic lesions and identify mutations in NDK in humans associated both with increased risk and protection from PD. We discovered that NDK lies at the intersection of proper ATP production, de novo synthesis of guanosine diphosphate and the activity of GTP cyclohydrolase I, a consumptive pathway needed to produce the bioactive metabolite tetrahydrobiopterin (BH_4_) required for mitochondrial function. Loss of NDK and impairment in guanosine nucleotide synthesis exacerbate synaptic dysfunction, while boosting the GTP consuming pathway promotes bioenergetics. Additionally, analysis of genetic data taken from over 64,000 PD affected individuals and 38,000 controls from diverse ancestries reveal that several genes lying at the intersection of guanosine nucleotide metabolism and bioenergetics pose a significant risk burden for PD.

---

The brain is a metabolically vulnerable organ and deficits in brain bioenergetic metabolism have long been considered early predictors of eventual neurodegeneration, particularly in PD^1,2^. We previously identified nerve terminal function as one of the loci of the brain’s metabolic vulnerability whereby synaptic vesicle (SV) recycling slows dramatically when local ATP production is curtailed^3^. We leveraged this finding to determine that a critical control point in nerve terminal ATP production is the glycolytic enzyme PGK1^4^, which had previously been identified as the primary off-target site of modulation by Terazosin (TZ), an FDA-approved α1 adrenergic receptor (A1R) antagonist used as treatment for benign prostate hyperplasia (BPH), that was shown to provide strong protection in numerous PD models^5^. Clinical use of TZ provided data for retrospective epidemiology that showed TZ dramatically lowers the likelihood of developing PD^6^ or dementia with Lewy Bodies^7^ compared to BPH patients taking an alternate A1R blocker. PGK1 is part of the PARK12 susceptibility locus^8^ mutations in PGK1 are associated with very early onset of the disease^9^. Overexpression of PGK1 in rodent mid-brain offered strong protection against substantia-nigral dopamine neuron degeneration^4^, while boosting ATP production via PGK1 could reverse the synaptic defect associated with PARK20^4^. In addition to ATP, a second purine metabolite, guanosine, is also a common currency for controlling numerous subcellular processes via the hydrolysis of guanosine triphosphate (GTP). Mutations in genes encoding two different enzymes in the pathway that converts GTP into the metabolite tetrahydrobiopterin (BH_4_) have been strongly linked to PD in humans^10–13^, while boosting GCH1 activity or increasing BH_4_ levels is strongly protective in murine PD models^14^. In the cytoplasm, the only pathway to synthesize GTP is via the enzyme NDK, which assembles as a hexamer of NDK1 and NDK2^15^ and transfers a phosphate from ATP onto GDP, suggesting a possible link between bioenergetic sensitivity and perturbations in guanosine nucleotide metabolism.

We leveraged large-scale genotyping and sequencing data release 11 of the Global Parkinson’s Genetics Program (GP2; https://gp2.org/) (https://doi.org/10.5281/zenodo.17753486) to determine if enzymes in this pathway might be linked to PD. This release comprises 103,786 genotyped participants (including 46,327 PD cases and 28,857 controls), as well as 38,266 whole-genome sequencing (WGS) participants (including 18,219 PD cases and 9,172 controls). Genotyping was performed using the NeuroBooster Aarray (NBA)^16^ that contains over 1.9 million variants, including approximately 10,000 custom content variants designed to improve imputation accuracy across diverse ancestries, as well as WGS to identify rare variants. Gene-level association analyses evaluated the contribution of variants in NME1 and NME2 to PD risk (NME1/2 are the human homologues of murine NDK1/2). Common variants (MAF ≥1%) were tested using NBA genotyping imputed and WGS data by logistic regression, whereas rare variants (MAF <1%) were analyzed in WGS data using SKAT and SKAT-O following grouping by allele frequency and functional annotation. In the NBA dataset, ancestry-stratified single-variant analyses identified a significant association for NME2 in the Central Asian group (Fig. 1A, Table S1) while SKAT-O and SKAT identified significant associations for NME1 and NME2 (Table S2 - 3) in the European, African, East Asian and central Asian groups.

**Figure 1.**
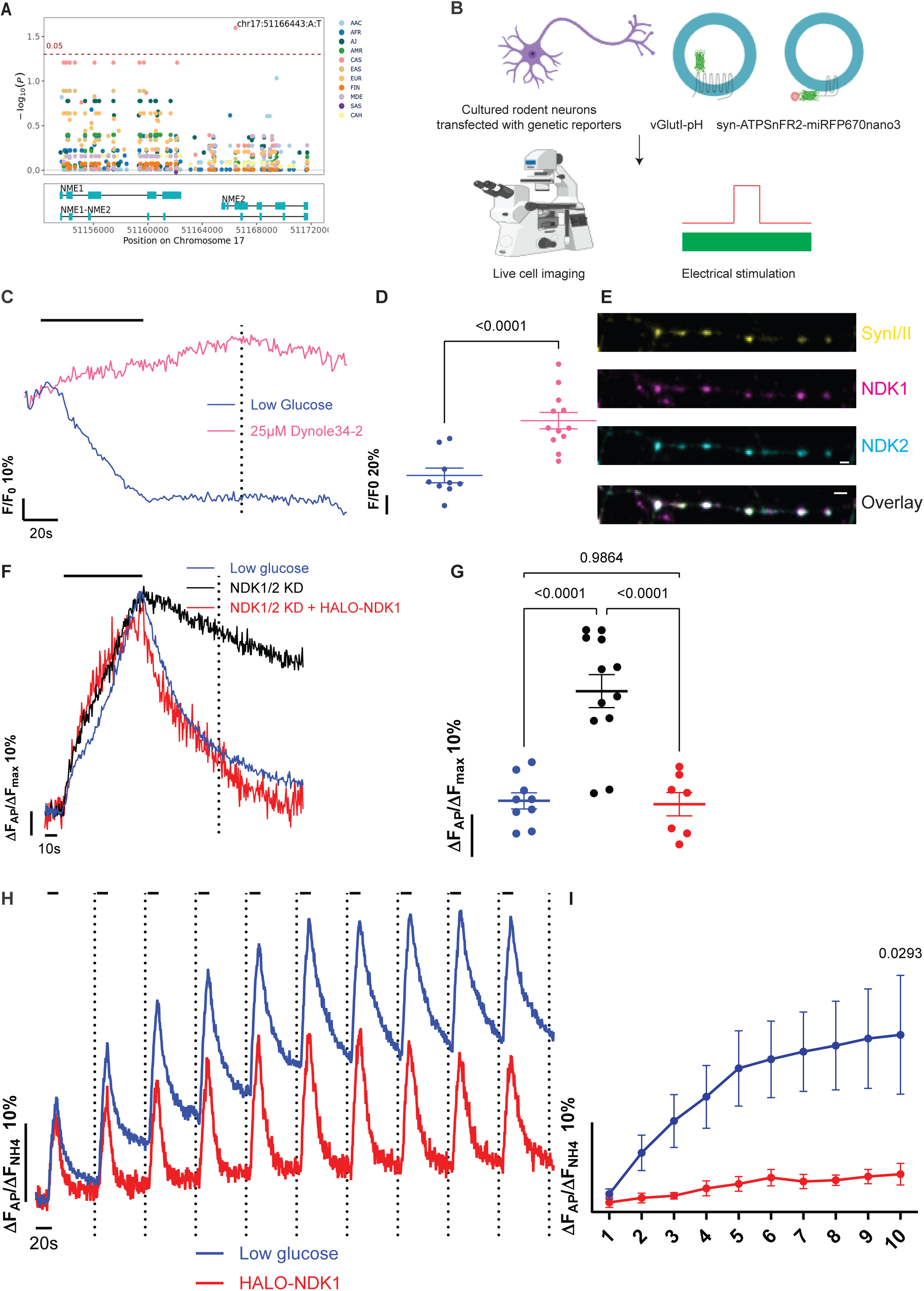
NDK1/2 are necessary GTP effectors for physiological synaptic transmission. (A) Regional plot of the NME1/NME2 locus. Association results are shown for the NME1 and NME2 region (chr17:51,153,559–51,171,749) across all ancestry groups. Analyses were performed using an adjusted regression model with a minor allele frequency (MAF) filter of >1%. The y-axis displays –log₁₀(p) values after Benjamini–Hochberg false discovery rate (FDR-BH) correction, while the x-axis corresponds to genomic positions. Significant variants are annotated by their IDs, with the horizontal red dashed line marking the 0.05 significance threshold. The lower panel depicts exonic regions as blue boxes. Cellular live-cell imaging schematic. In brief, primary rodent neurons were cultured, transfected with optical genetic reporters and imaged live, while electrically stimulated to trigger synaptic transmission. (C) Average syn-ATPSnFR2.0-miRFP670nano3 traces in 0.1 mM glucose (blue) or in acute presence of 25 μΜ Dynole34-2 (pink) normalized to baseline, where neurons were challenged with 600 APs at 10 Hz (black bar). (D) The remaining ATP values were quantified 60 s post stimulation (black dotted line in C) mean ± SEM, N=9 and 12 respectively, unpaired t-test. Blocking the GTPase activity of Dynamin I during neuronal activity, sustained synaptic ATP levels. (E) Immunostaining of primary neurons against NDK1 (magenta), NDK2 (cyan) and a nerve terminal marker, syn I/II (yellow), shows the synaptic localization of NDK1/2, scale bar 2 μm. (F) Average vGlutI- pHl traces in 0.1 mM glucose (blue), NDK1/2 KD (black) and rescue with HALO-NDK1 (red) normalized to peak height, where neurons were challenged with 600 APs at 10 Hz (black bar). (G) The remaining vGlutI-pHl fluorescence values, a reporter of SV endocytosis, were quantified 60 s post stimulation (black dotted line in F) mean ± SEM, N=9, 12 and 7 respectively, ordinary 1-way ANOVA. NDK1/2 are necessary for physiological synaptic transmission under hypometabolic conditions. (H) Average vGlutI- pHl traces in 0.1 mM glucose (blue) and HALO-NDK1 (red) normalized to maximum fluorescence revealed by perfusion of 50 mM NH_4_Cl, where neurons were challenged with 100 APs at 10 Hz every 1 min (black bars). (I) The remaining vGlutI-pHl fluorescence values were quantified 55 s post stimulation (black dotted lines in I), mean ± SEM, N=13 and 9 respectively, 2-way ANOVA. NDK1 overexpression safeguards synaptic transmission under hypometabolic conditions.

NDK (the sole fly gene encoding this enzyme) had previously been identified in a genetic screen in drosophila, that when mutated, worsens the phenotype associated with mutations in dynamin^17^, the central mechanochemical GTPase in cells that carries out membrane fission, including during SV recycling at nerve terminals^18^. We previously showed that genetic ablation of the major neuronal dynamin isoforms leads to a severe block of SV endocytosis^19,20^, comparable to that observed during metabolic compromise^21^. We had determined that one of the likely metabolic susceptibility loci to acute fuel deprivation in the brain are nerve terminals as they undergo complete arrest in their ability to recycle synaptic vesicles (SVs) when local ATP production is curtailed^3^. Given that GTP is primarily derived from ATP we sought to determine if acutely blocking dynamin function in primary neurons (Fig. 1B) would suppress ATP consumption during electrical activity in nerve terminals. Under low fuel conditions, acute action potential (AP) firing leads to a ∼ 40% drop in nerve terminal ATP that is completely suppressed with application of small molecule dynamin inhibitor^22^ that blocks SV endocytosis (Fig. 1C, D, S1A, B). This data strongly implies that SV endocytosis is necessary to trigger the major activity-driven ATP consumption at nerve terminals. As expected NDK1 and NDK2 are both found at nerve terminals (Fig. 1E) and a 50% reduction in NDK1/2 expression (Fig. S2A - C) impairs SV endocytosis measured with a pHluorin-tagged vesicular glutamate transporter (vGlutI-pHl) when glucose levels are restricted (Fig. 1F, G) but not under control conditions (Fig. S1E, F). Re-expression of an shRNA-resistant HALO-tagged NDK1 fully reversed the endocytic block (Fig. 1F, G). Similar results were obtained using acute pharmacological inhibition of NDK (Fig. S2G, H). As a second approach to establish that NDK1 phenotypes are metabolic, we activated PGK1 using TZ under restrictive glucose concentration, a mechanism that boosts synaptic ATP dynamics^4^, which rescued the NDK1 KD phenotype (Fig. S2I, J). In agreement with previous findings that metabolic compromise specifically affects SV endocytosis, exocytosis remained unaffected by these manipulations (Fig. S1D). The selective sensitivity of partial loss of NDK to metabolic compromise supports the idea that the rate of GTP production is determined by a combination of both ATP and NDK abundance and suggests that it may be possible to suppress slowed SV cycling when ATP production is curtailed by increasing NDK expression. To test this idea, we made use of a previously developed synaptic endurance test^4^, whereby nerve terminals are challenged with bursts of electrical activity at minute intervals in neurons expressing vGlutI-pHl to track SV recycling. Under low glucose conditions, this synaptic endurance fails when extracellular glucose is low (Fig. 1H) as successive rounds of stimulation lead to a gradual slowing of the endocytic retrieval/reacidification process and a gradual increase in the post stimulus fluorescence (Fig. 1I). We found the expression of HALO-tagged NDK1 was able to suppress this hypometabolic phenotype, allowing virtually normal SV cycling during 10 rounds of stimulation (Fig. 1I, J). These data collectively establish that NDKs are a metabolically sensitive control point in synaptic transmission.

Given that the human NDK genes (NME1 and NME2) are associated with increased PD risk (Fig. 1A, Table S1 - 3), we sought to assess whether NDKs might offer protection to acute degeneration of dopaminergic neurons in-vivo. AAV-mediated expression of mRuby-NDK1 in mouse midbrain (Fig. 2A), led to a significant increase in survival of animals that had received a unilateral 6-hydroxy-dopamine (6-OHDA) injection into the medial forebrain bundle compared to animals that were previously injected with an AAV driving mRuby (Fig S3A). Quantification of Tyrosine Hydroxylase (TH) positive dopaminergic neurons in the SNc, showed that NDK1 expression significantly protected these neurons from 6-OHDA mediated degeneration (Fig. 2B, C, S3B).

**Figure 2.**
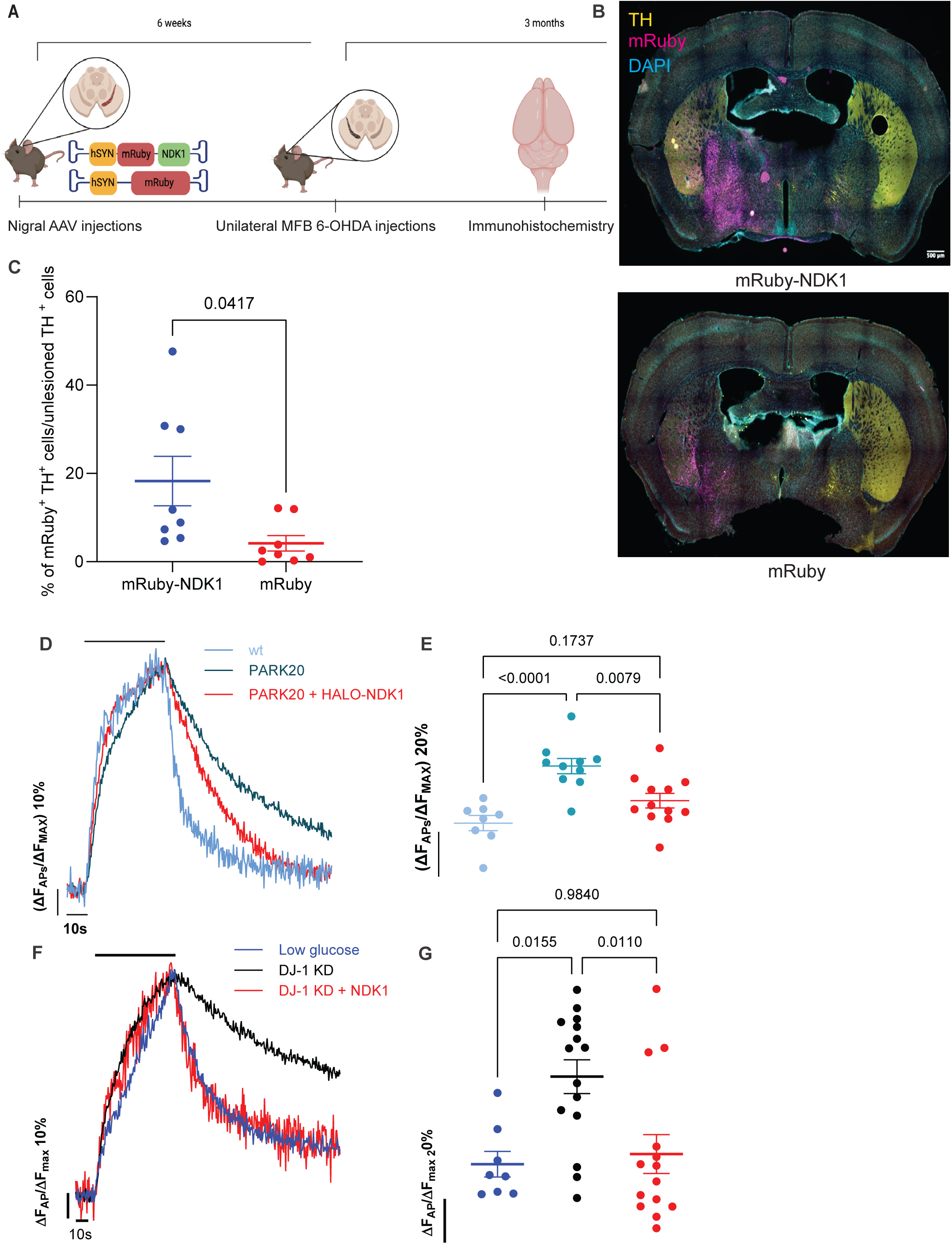
NDK1 protects against PD-associated synaptic dysfunction. (A) In-vivo 6-OHDA assay schematic. In brief, mice were injected with either mRuby-NDK1 or mRuby control AAV and 6 weeks later injected unilaterally with 6-OHDA. Their brains were subsequently immunostained with a-TH (yellow), DAPI for nuclear staining (cyan), while mRuby was conserved from the AAV injections (magenta). (B) Representative whole brain images of mRuby-NDK1 (top) or mRuby (bottom) animals show the specific protection of TH^+^ midbrain dopaminergic neurons by the NDK1 expression, scale bar 50 μm. (C) Quantification of mRuby and TH double positive neurons from mRuby-NDK1 (red) and mRuby (blue) animals, normalized to the non-lesioned side TH^+^, mean ± SEM, N=8 and 8 respectively, unpaired t-test. (D) Average vGlutI-pHl traces in 5 mM glucose of wt (light blue), PARK20 littermates (dark blue) and PARK20 + HALO-NDK1 (red) normalized to peak height, where neurons were challenged with 600 APs at 10 Hz (black bar). (E) The remaining vGlutI-pHl fluorescence values were quantified 60 s post stimulation, mean ± SEM, N=8, 10 and 12 respectively, 2-way ANOVA. NDK1 restored defective synaptic transmission, caused by PARK20 mutation. (F) Average vGlutI-pHl traces in 0.1 mM glucose (blue), DJ-1 KD, mimicking loss of function PARK7 (black) and DJ-1 KD + HALO-NDK1 (red) normalized to peak height, where neurons were challenged with 600 APs at 10 Hz (black bar). (G) The remaining vGlutI-pHl fluorescence values were quantified 60 s post stimulation, mean ± SEM, N=8, 15 and 14 respectively, 2-way ANOVA. NDK1 restored defective synaptic transmission, caused by absence of DJ-1/PARK7.

We proceeded to interrogate the functional consequences of NDK1 in two genetic models of PD. First, we used PARK20 primary mouse neurons bearing the R258Q mutation in Synaptojanin I, which is characterized by defective SV recycling^23^. NDK1 OE in these neurons restored physiological synaptic function (Fig. 2D, E) and had no effect in wild type (wt) litter-mate neurons (Fig. S3C, D). The second PD model we employed is the PARK7 model, characterized by loss-of-function of the chaperone DJ-1 and defective SV recycling^4,24^. We previously showed that DJ-1 is a likely PGK1 chaperone, as loss of DJ-1 leads to a profound inability to adequately produce ATP in nerve terminals following activity, and glycolytic activation by either PGK1 expression or TZ application fails in the absence of this protein^4^. In contrast, we found that directly promoting GTP production, by boosting NDK1 expression, restored defective synaptic function (Fig. 2F, G) when DJ-1 expression is suppressed. These data collectively suggest that GTP homeostasis is a previously unknown pathophysiological synaptic pathway implicated in PD and can be successfully exploited both in-vivo and in genetic models of PD to restore defective dopaminergic transmission.

Our data indicates that both NDK and ATP abundance are critical in determining if sufficient GTP is produced to support SV recycling in both healthy neurons and those harboring PD-driving mutations. The third component required for GTP production is an adequate supply of GDP. Although produced by GTP hydrolysis, we sought to determine whether synapses also relied on de-novo production of GDP which synthesizes GTP from inosine-5′-monophosphate (IMP) via a sequence of enzymatic steps by IMP dehydrogenase (IMPDH) to produce xanthine-monophosphate (XMP), then converted to guanosine monophosphate (GMP) by guanosine monophosphate dehydrogenase (GMPS) and phosphorylated by guanosine kinase I (GUK1) to produce GDP (Fig 3A). IMPDH1 and IMPDH2 are both present in nerve terminals (Fig. 3B) and strikingly, short-term pharmacological inhibition of these enzymes for 2 h, using two different classes of drugs, arrested SV endocytosis (Fig. 3E, F, S4G, H) under restricted glucose conditions. These data indicate that nerve terminals rely acutely on constitutive de-novo GDP synthesis. Similarly, specific KD of each isoform (Fig. S4A - D) slowed SV endocytosis (Fig. S4E, F).

**Figure 3.**
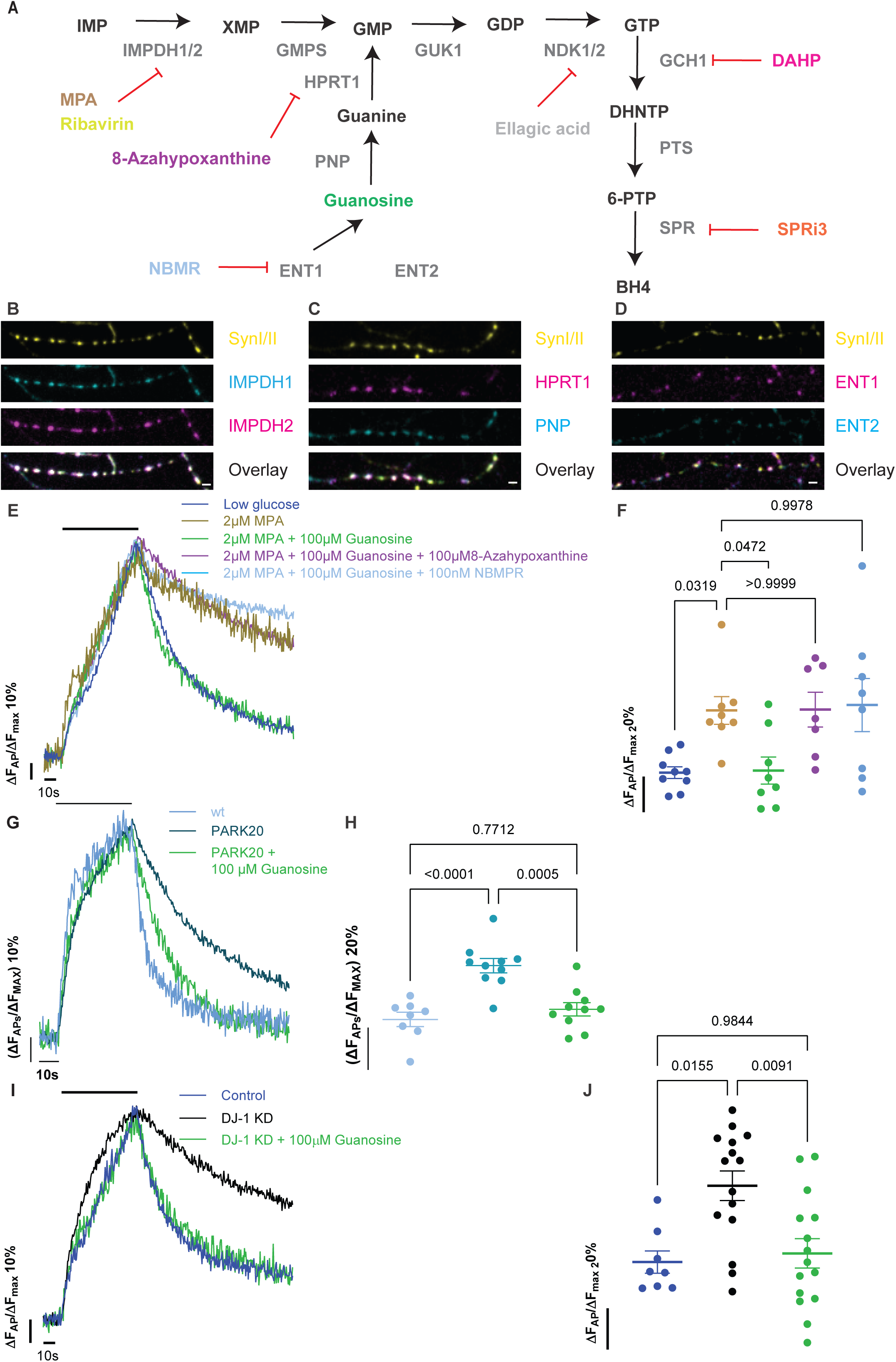
Guanosine nucleotide biosynthetic pathways are crucial for synaptic transmission. (A) Outline of both the de-novo, salvage guanosine nucleotide and BH_4_ biosynthetic pathways, along with the inhibitors used for some of these enzymes. Immunostaining of primary neurons against (B) IMPDH1 (cyan), IMPDH2 (magenta), (C) HPRT1 (magenta), PNP (cyan), (D) ENT1 (magenta), ENT2 (cyan) and syn I/II (yellow) shows the synaptic localization of these enzymes, scale bar 2 μm. (E) Average vGlutI-pHl traces in 0.1 mM glucose (blue), 2 μM MPA (brown), 2 μM MPA + 100 μΜ Guanosine (green), 2 μM MPA + 100 μΜ Guanosine + 100 μM 8-Azahypoxanthine (purple) and 2 μM MPA + 100 μΜ Guanosine + 100 nM NBMPR (light blue) all incubated for 2 h, normalized to peak height, where neurons were challenged with 600 APs at 10 Hz (black bar). (F) The remaining vGlutI-pHl fluorescence values were quantified 60 s post stimulation, mean ± SEM, N=9, 8, 8, 7 and 8 respectively, 2-way ANOVA. Blocking the de-novo guanosine nucleotide synthetic pathway impacts synaptic transmission and can be suppressed by supplementation of Guanosine, which enters cells via ENT1 and acts via HPRT1 of the salvage pathway. (G) Average vGlutI-pHl traces in 5 mM glucose of wt (light blue), PARK20 littermates (dark blue) and PARK20 + 100 μΜ Guanosine incubated for 2 h (green) normalized to peak height, where neurons were challenged with 600 APs at 10 Hz (black bar). (H) The remaining vGlutI-pHl fluorescence values were quantified 60 s post stimulation, mean ± SEM, N=8, 10 and 10 respectively, 2-way ANOVA. Guanosine restored defective synaptic transmission, caused by PARK20 mutation. (I) Average vGlutI-pHl traces in 0.1 mM glucose (blue), DJ-1 KD, mimicking loss of function PARK7 (black) and DJ-1 KD + 100 μΜ Guanosine incubated for 2 h (green) normalized to peak height, where neurons were challenged with 600 APs at 10 Hz (black bar). (J) The remaining vGlutI-pHl fluorescence values were quantified 60 s post stimulation, mean ± SEM, N=8, 15 and 15 respectively, 2-way ANOVA. Guanosine restored defective synaptic transmission, caused by absence of DJ-1/PARK7.

In addition to de-novo synthesis of GDP, cells can also rely on the guanosine salvage pathway, whereby guanosine (either taken up from the extracellular space by the equilibrative Nucleoside Transporters (ENT1/2) or salvaged from other metabolites) is converted to guanine by purine nucleoside phosphorylase (PNP) and then to GMP via Hypoxanthine phosphoribosyltransferase 1 (HPRT1). Remarkably all three of these enzymes are enriched in nerve terminals (Fig. 3C). The presence of this pathway at nerve terminals suggests that it may be possible to directly drive local GDP production via acute addition of extracellular guanosine. To test this idea, we simultaneously added extracellular guanosine while inhibiting IMPDH1/2. These experiments demonstrated that the salvage pathway, provided sufficient extracellular guanosine is available, can bypass the de-novo synthesis pathway at nerve terminals (Fig. 3E, F, S4G, H). This guanosine rescue was fully inhibited however, if either HPRT1 or ENT1 were inhibited (Fig. 3D - F). Similarly incubating PARK20 neurons or neurons expressing a DJ-1 shRNA (PARK7) rescued the slow SV recycling phenotype when preincubated with extracellular guanosine (Fig 3G - J).

The unexpected sensitivity of synaptic transmission to acute perturbation in GDP synthesis (Fig. 3E, F) suggests that GTP is locally consumed in a pathway that does not produce GDP. A strong candidate for this is GTP cyclohydrolase 1 (GCH1), the first of three enzymes, followed by 6-pyruvoyltetrahydropterin synthase, PTS, and sepiatperin reductase, SPR, required for tetrahydrobiopterin (BH_4_) biosynthesis (Fig. 3A). Importantly, mutations in GCH1 have recently been linked to PD^10,13,25^, while SPR, is considered the key gene conferring PD risk in the PARK3 locus^11^. Both GCH1 and SPR are present in nerve terminals (Fig. 4A) suggesting they play a local presynaptic role in guanosine metabolism. Although BH_4_ is a necessary co-factor for several amino acid hydroxylases^26^, recent work has demonstrated that this metabolite is also necessary for mitochondrial function^14,27^. We tested this hypothesis that BH_4_ production is necessary for axonal bioenergetic support by determining if blocking its production would exacerbate synapse function under metabolic stress. Both shRNA-mediated suppression (Fig. S5A, B) and acute pharmacological inhibition (by 2,4-diamino-6-hydroxypyrimidine, DAHP) of GCH1 slowed SV recycling under restrictive glucose conditions (Fig. 4B, C) while the former would be rescued by reintroduction of a shRNA resistant HALO tagged GCH1 (Fig. 4B, C). Similar results were obtained using an SPR inhibitor, SPRi3 (Fig. S5C, D). To further test the idea that BH_4_ production directly controls local synaptic bioenergetics, we examined the dynamics of presynaptic ATP during and after acute electrical stimulation under restrictive glucose conditions in neurons that were either preincubated with BH_4_ or over-expressing GHC1. These experiments showed that boosting BH_4_ availability profoundly suppressed activity driven loss of ATP during stimulation and accelerated recovery of ATP following the stimulus period (Fig. 4D, E). Similar experiments carried out in the presence of oligomycin (Fig. S5E, F), showed that this GCH1/BH_4_ mediated boosting of synaptic ATP productions was mediated by mitochondria, consistent with previous observations^27^. The ability of GCH1 enhancement to boost local ATP production suggests that this approach should also protect nerve terminal function under hypoglycemic conditions. GCH1 OE indeed protected synaptic transmission from failure in our synaptic endurance assay (Fig. 4F, G). Our data collectively suggests that the BH_4_ biosynthesis pathway is an important pathophysiological pathway in PD, linking both GTP and ATP pathways. These results are consistent with the idea that GTP production is being balanced by a consumption pathway that is also necessary to sustain axonal bioenergetics.

**Figure 4.**
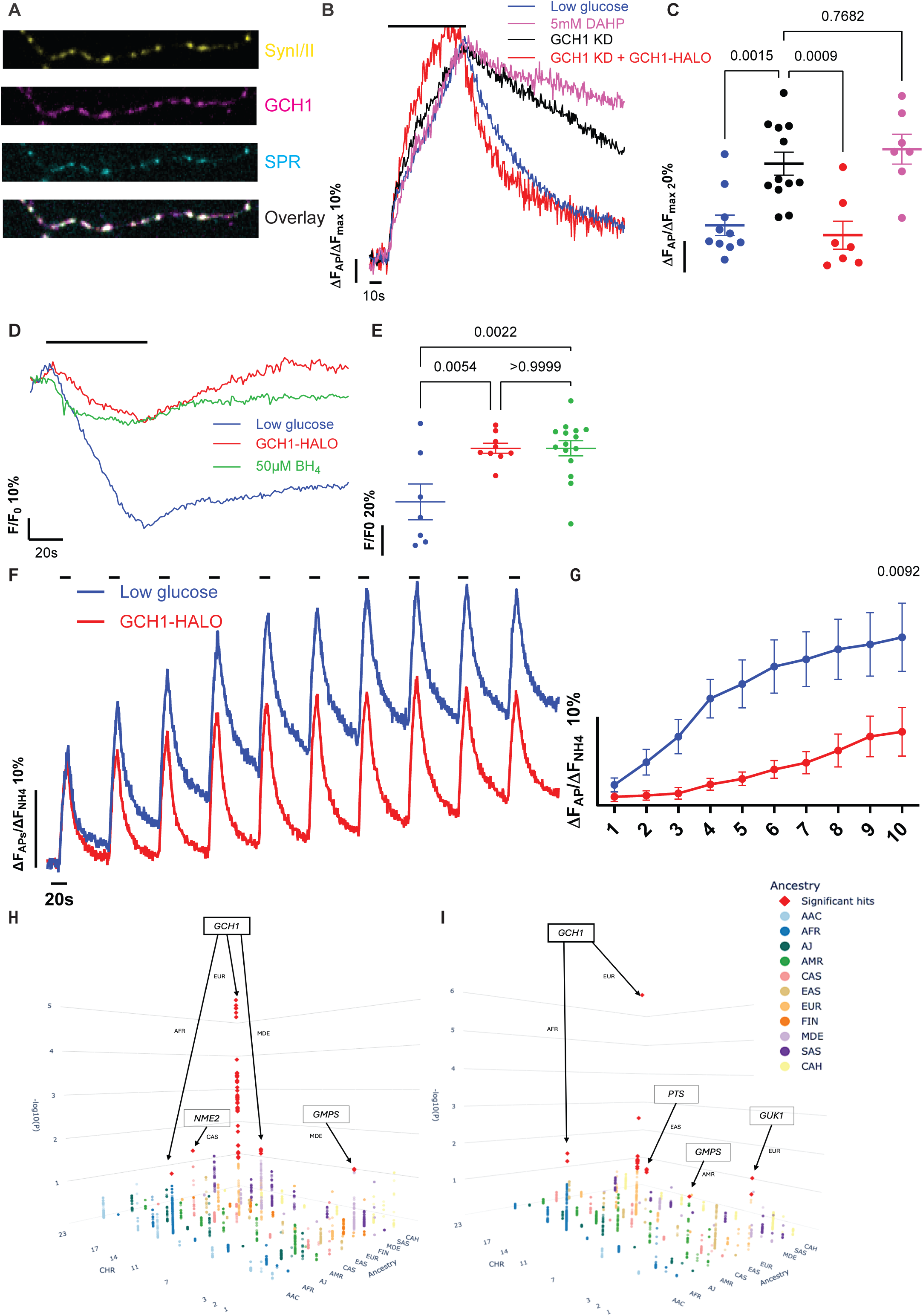
The BH_4_ biosynthetic pathway is crucial for synaptic transmission by locally providing ATP. (A) Immunostaining of primary neurons against GCH1 (magenta), SPR (cyan) and syn I/II (yellow) shows the synaptic localization of GCH1 and SPR, scale bar 2 μm. (B) Average vGlutI-pHl traces in 0.1 mM glucose (blue), GCH1 KD (black), GCH1-HALO rescue (red) and 5 mM DAHP incubated ON (magenta) normalized to peak height, where neurons were challenged with 600 APs at 10 Hz (black bar). (C) The remaining vGlutI-pHl fluorescence values were quantified 60 s post stimulation, mean ± SEM, N=10, 12, 7 and 7 respectively, 2-way ANOVA. Blocking the BH_4_ biosynthetic pathway impacts synaptic transmission. (D) Average syn-ATPSnFR2.0-miRFP670nano3 traces in 0.1 mM glucose (blue), GCH1-HALO (red) and 50 μΜ BH_4_ incubated ON (green) normalized to baseline, where neurons were challenged with 600 APs at 10 Hz (black bar). (E) The remaining ATP values were quantified at the end of the stimulation, mean ± SEM, N=7, 9 and 15 respectively, ordinary 1-way ANOVA. Upregulation of BH_4_ promotes ATP production. (F) Average vGlutI-pHl traces in 0.1 mM glucose (blue) and GCH1-HALO (red) normalized to maximum fluorescence revealed by perfusion of 50 mM NH_4_Cl, where neurons were challenged with 100 APs at 10 Hz every 1 min (black bars). (G) The remaining vGlutI-pHl fluorescence values were quantified 55 s post stimulation, mean ± SEM, N=16 and 15 respectively, 2-way ANOVA. Genome-wide association signals are displayed for SPR, PTS, PNP, NME1, NME2, IMPDH1, IMPDH2, HPRT1, GUK1, GMPS, and GCH1 across ancestry groups using (H) NBA genotyping imputed data and (I) whole-genome sequencing (WGS) data. Analyses were conducted with generalized linear models (GLM), adjusting for age, sex, and the first five genetic principal components. Each point represents a tested variant, with chromosome position, –log₁₀(p), and ancestry groups mapped to the three axes. Variants significant after FDR-BH correction are highlighted as red diamonds. Abbreviations: AAC, African Admixed; AFR, African; AJ, Ashkenazi Jewish; AMR, American Admixed; CAS, Central Asian; EAS, East Asian; EUR, European; FIN, Finnish; MDE, Middle Eastern; SAS, South Asian; CAH, Complex Admixture. Abbreviations: AAC, African Admixed; AFR, African; AJ, Ashkenazi Jewish; AMR, American Admixed; CAS, Central Asian; EAS, East Asian; EUR, European; FIN, Finnish; MDE, Middle Eastern; SAS, South Asian; CAH; Complex Admixture.

We leveraged the same large-scale genotyping and sequencing data release 11 of the Global Parkinson’s Genetics Program used to identify NME2 as a PD risk gene, to determine whether mutations in this suite of guanosine metabolism enzymes are associated with PD risk. Gene-based rare variant analyses using SKAT identified significant nominal associations for GCH1, GUK1, HPRT1, PNP, and PTS across multiple ancestry groups, and variant categories (Table S1 - 3). GCH1 showed associations in both African and European groups under the rare variant category (MAF<1%) as well as in the African group under the potentially functional category. SKAT-O showed a similar overall pattern of associations while identifying significant associations for SPR, and additional ancestry-specific signals for GCH1 and GUK1 (Fig. 4H, I). This analysis identified multiple mutations in both the de-novo synthesis and salvage pathway enzymes that can both increase and decrease PD risk (Fig. S5G, Table S1 – 3).

## Discussion

Using a mix of functional imaging in primary neurons, in-vivo experiments in rodents and large human genome dataset analysis from novel ancestries, we show that deficits in guanosine nucleotide metabolism constitutes a previously unknown pathogenic mechanism in PD that is closely intertwined with cellular bioenergetics. Of special importance are the enzymes NDK1/2, as they bridge ATP and GTP metabolism (Fig. S5H) and show how two distinct PD susceptibility pathways are closely linked. The findings of their outsized importance in synaptic transmission both in-vivo and in-vitro, also elucidate a previously open question in the field which is why a bioenergetic failure arrests SV recycling. Our data show that this metabolic vulnerability arises from the close connection between ATP and GTP production. That GTP itself has critical importance is supported by the fact that both overexpression of NDK or simple supplementation with extracellular guanosine can offer protection against metabolic lesions or genetic mutations linked to PD. This experimental evidence is fully supported by human genomic data, since several metabolic enzymes we identified are associated both with decreased and increased PD risk respectively.

Additionally, there is evidence that guanosine can play a neuroprotective role in neurodegeneration^28,29^, although the mechanistic basis for this had not been firmly established. We now establish that guanosine acts intracellularly as a physiological substrate of PNP and HPRT1 in the guanosine salvage pathway to promote protection of synapse function during a metabolic compromise. Our findings also pave the way for future studies on HPRT1, a protein responsible for Lesch-Nyhan syndrome, suggesting a critical GTP function in early brain development.

One surprising finding was that nerve terminals became intolerant of fuel restriction only two hours after blocking the de-novo synthesis pathway (Fig. 3), implying that over this time scale the pool of GDP/GTP is being depleted. We discovered that in addition to having enrichment in the synthetic pathways to generate GDP and GTP (Fig. 1E, 3B - D), nerve terminals are also enriched in enzymes that drive GTP consumption (Fig. 4A), providing a potential explanation for the relatively acute sensitivity to inhibiting de novo GDP synthesis. Although loss of consumption of GTP by GCH1 would normally sensitize synapses to bioenergetic stress owing to the need for greater ATP concentration to compensate in driving NDK, we showed that the metabolite BH_4_ produced by this pathway acts to boost mitochondrial function (Fig. 4D) in agreement with findings in T cells^27^. Given our findings that BH_4_, in addition to its known roles as a key co-factor for several amino acid hydroxylases, plays an essential role in axonal bioenergetics, the work we show here suggests that the local guanosine metabolism therefore likely plays a dual role with regards to PD susceptibility. Deleterious mutations in genes that impact GDP or GTP production as well as those that are required to produce BH_4_ will impact both dopamine synthesis, as tyrosine hydroxylase depends on BH4, as well as bioenergetics. Conditions that lead to deficits in ATP production will in turn create deficits in GTP production, because it would lead to a deficit in BH_4_ production would then exacerbate bioenergetics since it would impact mitochondria. The combination of cellular, in vivo mouse, and human genetic data all support the idea that the intersection of guanosine and adenosine metabolism is an important pathway in conferring risk in developing PD.

## Acknowledgments

We would like to thank members of the Ryan laboratory for discussions and L. Lavis for the gift of JF dyes and Andy Singleton for helping us integrate our findings with human genetic data. Funding: This research was supported in part by NIH grants NS036942 and NS11739 to T.A.R., a grant from JPB Foundation for Medical Research to M.S.K., Aligning Science Across Parkinson’s ASAP-000580 and through the Michael J. Fox Foundation for Parkinson’s Research (MJFF). Data (DOI XX/zenodo.xx,release x) used in the preparation of this article were obtained from the Global Parkinson’s Genetics Program (GP2). GP2 is funded by the Aligning Science Across Parkinson’s (ASAP) initiative and implemented by The Michael J. Fox Foundation for Parkinson’s Research (https://gp2.org).

## Author contributions

ACK: experimental design, data acquisition and manuscript preparation; SRU – experimental design and data acquisition; PR genetic data analysis; MI genetic data analysis; MTP genetic data analysis; MGK experimental design; TAR experimental design and manuscript preparation; GP2 genetic data acquisition.

## Competing interests

For the purpose of open access, we have applied a CC BY public copyright license to all Author Accepted Manuscripts arising from this submission. The authors declare that they have no other competing interests.

## Data and reagents availability

The data, code, protocols, and key lab materials used and generated in this study are listed in a Key Resource Table alongside their persistent identifiers at 10.5281/zenodo.21510483. All experimental protocols are collected at dx.doi.org/10.17504/protocols.io.n2bvj5r3bgk5/v1.

**Primary neuronal culture**

dx.doi.org/10.17504/protocols.io.ewov1qxr2gr2/v1

**Live-cell imaging**

dx.doi.org/10.17504/protocols.io.q26g7pn4qgwz/v1

**Immunocytochemistry**

dx.doi.org/10.17504/protocols.io.ewov19z5olr2/v1

**vGlutI-pH analysis**

dx.doi.org/10.17504/protocols.io.bp2l6275zgqe/v1

**ATPSnFR analysis**

dx.doi.org/10.17504/protocols.io.n92ld8wk9v5b/v1

**AAV craniotomy**

dx.doi.org/10.17504/protocols.io.5jyl829q7l2w/v1

**6-OHDA lesions**

dx.doi.org/10.17504/protocols.io.8epv5r9o4g1b/v1

**Immunohistochemistry**

dx.doi.org/10.17504/protocols.io.3byl4956jgo5/v1

---

**Figure S1.**
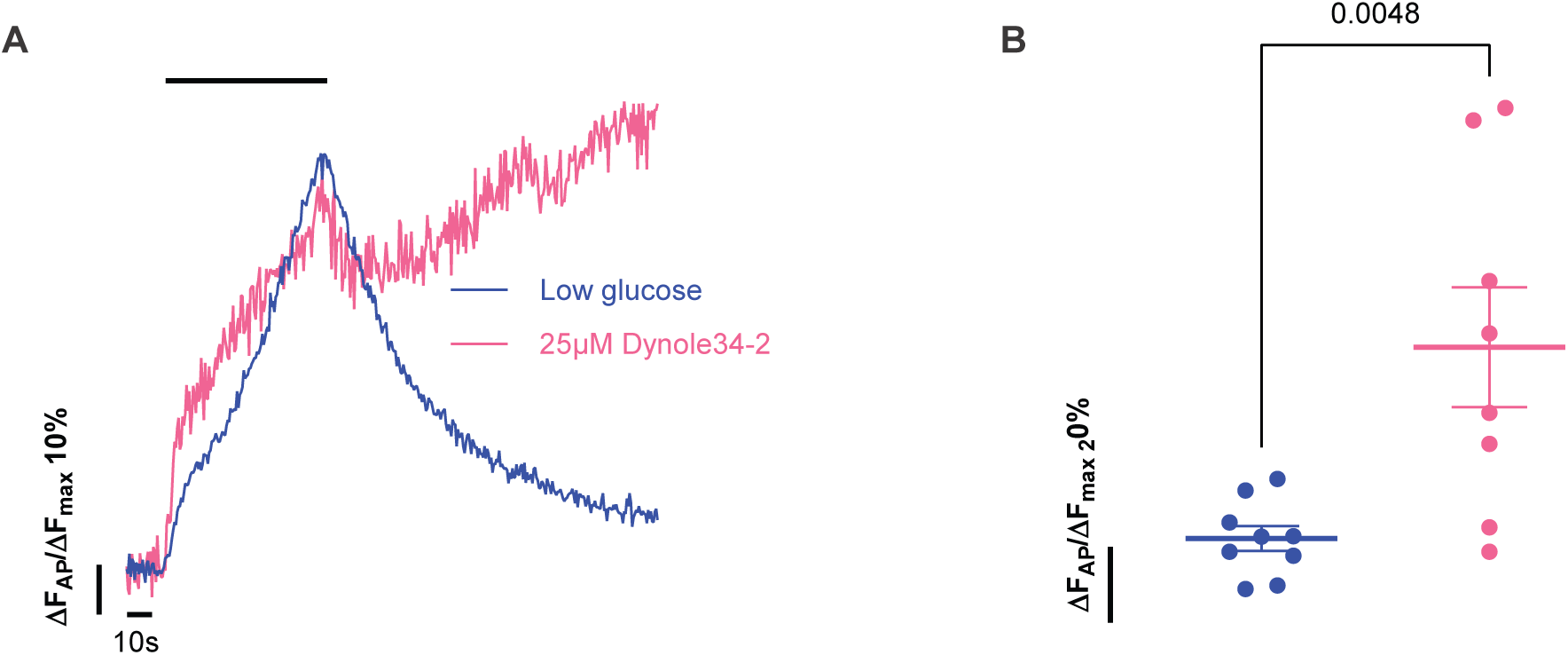
Dynole compounds validation. (A) Average vGlutI-pHl traces in 0.1 mM glucose (blue) and 25 μΜ Dynole 34-2 (pink) incubated acutely, normalized to peak height, where neurons were challenged with 600 APs at 10 Hz (black bar). (B) The remaining vGlutI-pHl fluorescence values were quantified 60 s post stimulation, mean ± SEM, N=9 and 8 respectively, unpaired t-test. Inhibition of the GTPase activity of Dynamin blocks SV endocytosis.

**Figure S2.**
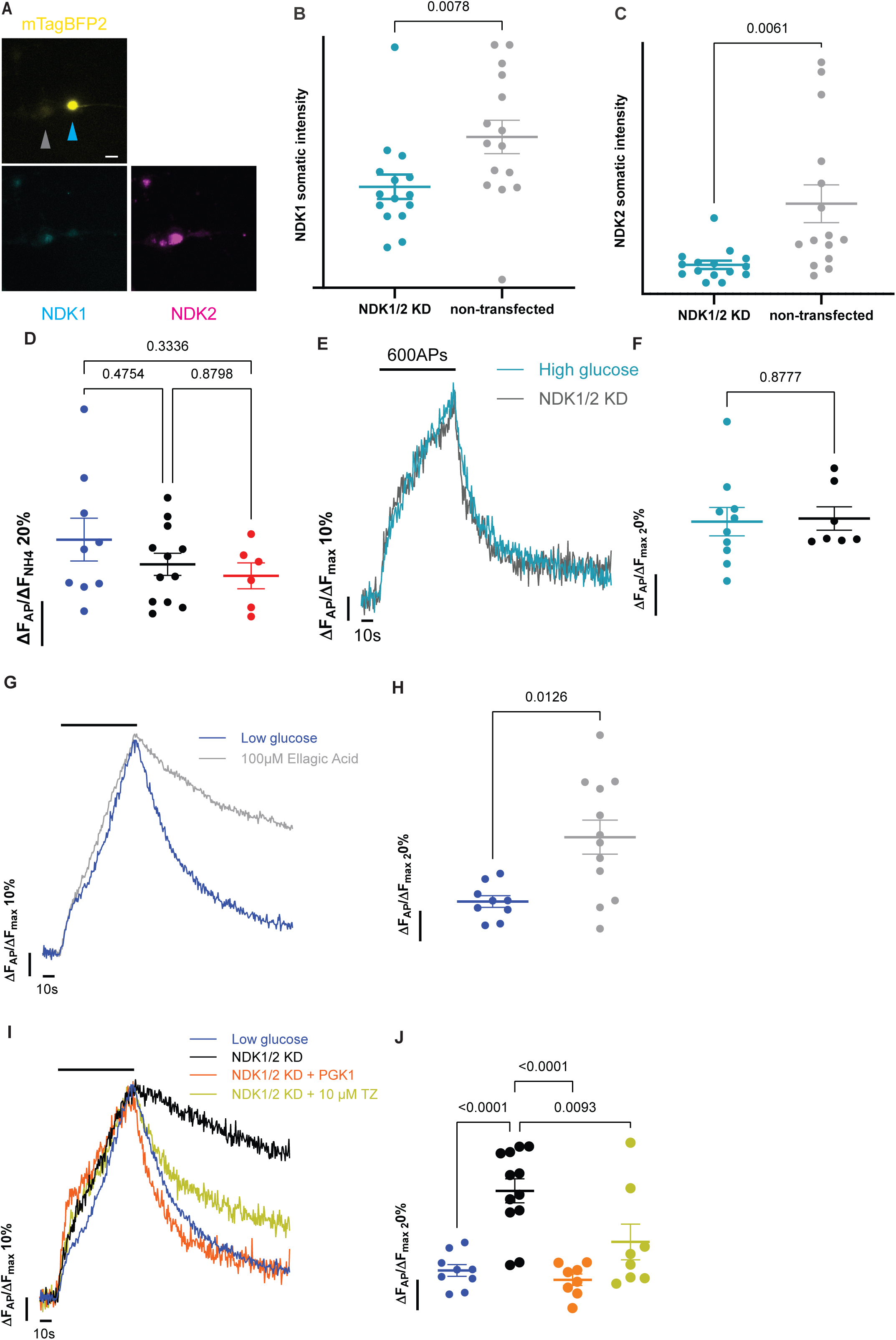
NDK1/2 KD can be suppressed by boosting ATP production. (A) Representative images of neurons expressing mTagBFP2 (yellow) along with NDK1/2 shRNA (cyan arrow) next to non-transfected (grey arrow) and immunostained against NDK1 (cyan) and NDK2 (magenta), scale bar 10 μm. Quantification of the KD efficiency of NDK1 (B) and NDK2 (C), mean ± SEM, paired t-test, N=15 for both (B) and (C). (D) Exocytic extend of the traces from Fig. 1G, ordinary 1-way ANOVA. (E) Average vGlutI- pHl traces in 5 mM glucose (teal) and NDK1/2 KD (grey) normalized to peak height, where neurons were challenged with 600 APs at 10 Hz (black bar). (F) The remaining vGlutI-pHl fluorescence values were quantified 60 s post stimulation, mean ± SEM, N=10 and 7 respectively, unpaired t-test. (G) Average vGlutI-pHl traces in 0.1 mM glucose (blue) and 100 μM Ellagic Acid (grey) added acutely normalized to peak height, where neurons were challenged with 600 APs at 10 Hz (black bar). (H) The remaining vGlutI-pHl fluorescence values were quantified 60 s post stimulation, mean ± SEM, N=9 and 12 respectively, unpaired t-test. Pharmacological inhibition of NDKs, mimics silencing of the enzymes. (I) Average vGlutI-pHl traces in 0.1 mM glucose (blue), NDK1/2 KD (black), NDK1/2 KD plus PGK1-HALO (orange) and NDK1/2 KD plus 10 μM Terazosin (yellow) normalized to peak height, where neurons were challenged with 600 APs at 10 Hz (black bar). (J) The remaining vGlutI-pHl fluorescence values were quantified 60 s post stimulation, mean ± SEM, N=9, 12, 12 and 8 respectively, ordinary 1-way ANOVA. Upregulation of ATP kinetics under hypometabolic conditions suppresses the NDK silencing.

**Figure S3.**
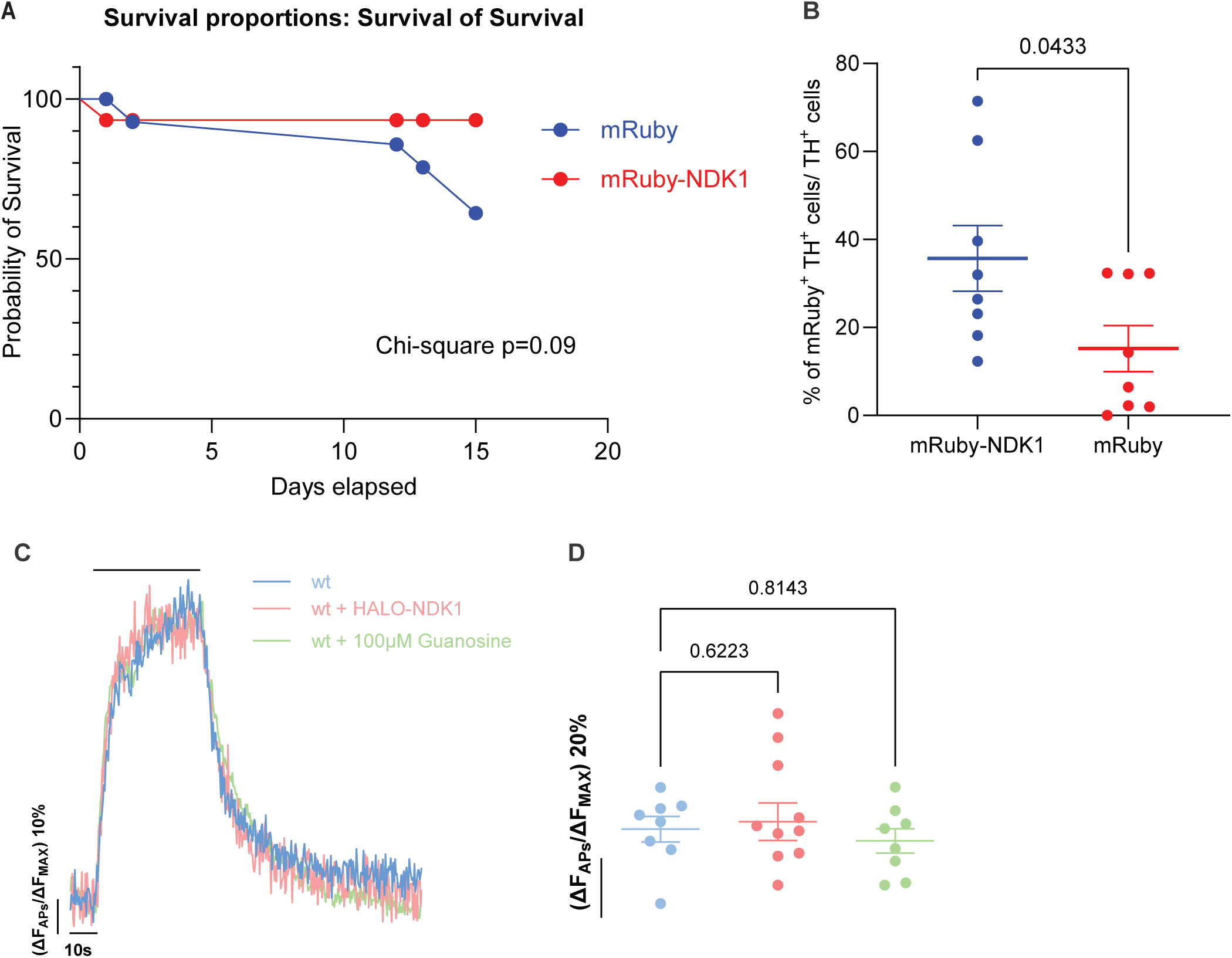
Guanosine can suppresses synaptic dysfunction caused by PARK mutations. (A) Survival analysis of the mRuby-NDK1 (red) and mRuby (blue) animals in Fig. 2A, following the 6-OHDA unilateral injection. NDK1 expression promoted animal survival. (B) Quantification of mRuby and TH double positive neurons from mRuby-NDK1 (red) and mRuby (blue) animals, normalized to the non-lesioned side TH+, mean ± SEM, N=8 and 8 respectively, unpaired t-test. (C) Average vGlutI-pHl traces in 5 mM glucose of wt (light blue), wt + HALO-NDK1 (light red) and wt + 100 μΜ Guanosine for 2 h (light green) normalized to peak height, where neurons were challenged with 600 APs at 10 Hz (black bar). (D) The remaining vGlutI-pHl fluorescence values were quantified 60 s post stimulation, mean ± SEM, N=8, 11 and 8 respectively, 2-way ANOVA. NDK1 and Guanosine had no effect in wt neurons.

**Figure S4.**
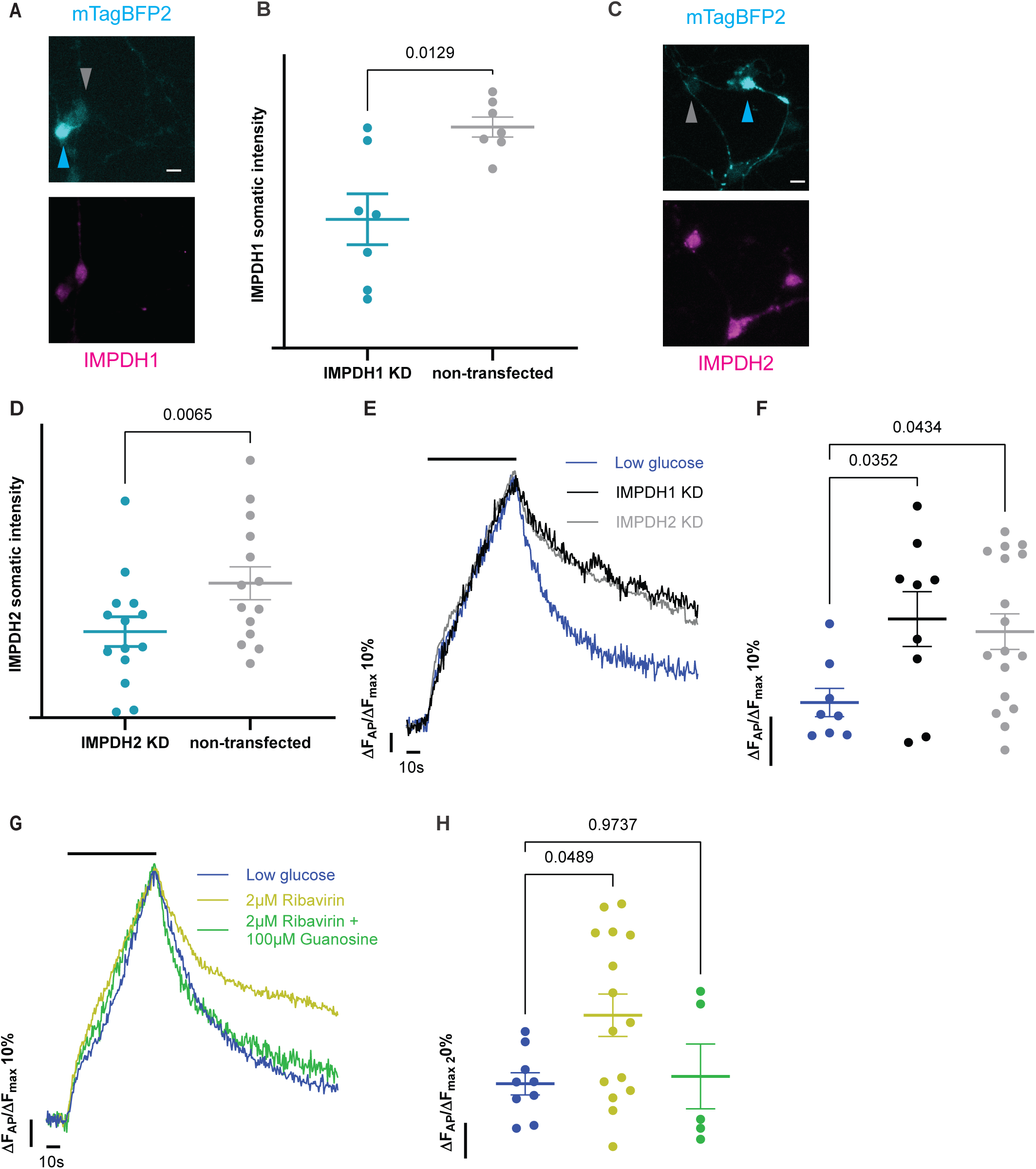
IMPDH1 and 2 KD validates the pharmacological inhibition. (A) Representative images of neurons expressing mTagBFP2 (cyan) along with IMPDH1 shRNA (cyan arrow) next to non-transfected (grey arrow) and immunostained against IMPDH1 (magenta), scale bar 10 μm. (B) Quantification of the KD efficiency of IMPDH1, mean ± SEM, N=7, paired t-test. (C) Representative images of neurons expressing mTagBFP2 (cyan) along with IMPDH2 shRNA (cyan arrow) next to non-transfected (grey arrow) and immunostained against IMPDH2 (magenta), scale bar 10 μm. (D) Quantification of the KD efficiency of IMPDH2, mean ± SEM, N=14, paired t-test. (E) Average vGlutI-pHl traces in 0.1 mM glucose (blue), IMPDH1 KD (black) and IMPDH2 (gray) normalized to peak height, where neurons were challenged with 600 APs at 10 Hz (black bar). (F) The remaining vGlutI-pHl fluorescence values were quantified 60 s post stimulation, mean ± SEM, N=8, 9 and 17 respectively, 2-way ANOVA. Blocking the rate limiting enzymes in guanosine nucleotide de-novo synthesis impacts synaptic transmission. (G) Average vGlutI-pHl traces in 0.1 mM glucose (blue), 2 μM Ribavirin (yellow) and 2 μM Ribavirin + 100 μM Guanosine incubated for 2 h (green) normalized to peak height, where neurons were challenged with 600 APs at 10 Hz (black bar). (H) The remaining vGlutI-pHl fluorescence values were quantified 60 s post stimulation, mean ± SEM, N=9, 15 and 5 respectively, 2-way ANOVA.

**Figure S5.**
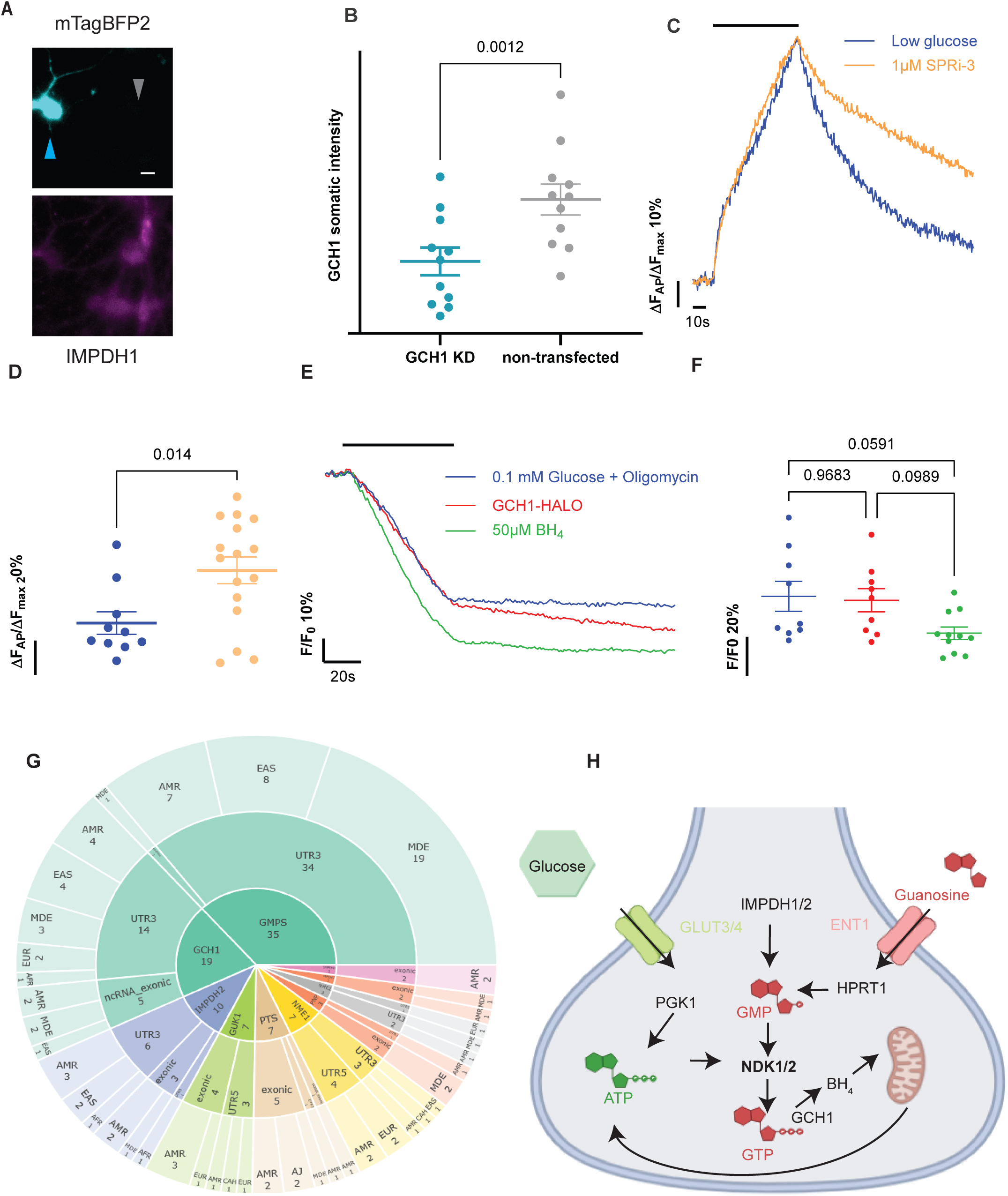
Mitochondria are necessary for the BH_4_ mediated upregulation of ATP production. (A) Representative images of neurons expressing mTagBFP2 (cyan) along with GCH1 shRNA (cyan arrow) next to non-transfected (grey arrow) and immunostained against GCH1 (magenta), scale bar 10 μm. (B) Quantification of the KD efficiency of GCH1, mean ± SEM, N=11, paired t-test. (C) Average vGlutI-pHl traces in 0.1 mM glucose (blue) and 1 μM SPRi3 incubated ON (orange) normalized to peak height, where neurons were challenged with 600 APs at 10 Hz (black bar). (D) The remaining vGlutI-pHl fluorescence values were quantified 60 s post stimulation, mean ± SEM, N=10 and 16 respectively, unpaired t-test. Blocking SPR, the BH_4_ synthetic enzyme, mimics the GCH1 inhibition. (E) Average syn-ATPSnFR2.0-miRFP670nano3 traces in 0.1 mM glucose + 2 μM Oligomycin (blue), GCH1-HALO (red) and 50 μΜ BH_4_ incubated ON (green) normalized to baseline, where neurons were challenged with 600 APs at 10 Hz (black bar). (F) The remaining ATP values were quantified at the end of the stimulation, mean ± SEM, N=9, 9 and 11 respectively, ordinary 1-way ANOVA. The BH_4_ mediated upregulation of ATP production required functional mitochondria. (G) Hierarchical sunburst visualization of significant single-gene associations derived from GP2 NBA and WGS datasets across 11 ancestry groups. Analyses were performed under unadjusted and covariate-adjusted models (age, sex, PC1–PC5). FDR–Benjamini–Hochberg correction was used. Intronic and intergenic variants were excluded to focus on coding and regulatory exonic signals. Segment size represents the number of significant associations per category. (H) Graphical summary and proposed mechanism. We had previously identified that PGK1 is the rate-limiting step in synaptic glycolysis, producing sufficient ATP to power synaptic transmission. In this study, we identify that guanosine nucleotide metabolic pathways have an outsized effect on synaptic transmission. In addition, we unravel NDKs as the enzymes connecting all these pathways and establish their implication in PD, using a pallet of cellular, in-vivo and human genetic approaches.

**Table S1.** Significant single-variant association results from logistic regression analyses adjusted for sex, age and the first five principal components (PCs). Abbreviations: A1, effect allele; CI, confidence interval; FDR-BH, Benjamini–Hochberg false discovery rate; Freq, frequency; HGVS, Human Genome Variation Society; NBA, NeuroBooster Array; OR, odds ratio; PCs, principal components; WGS, whole-genome

**Table S2.** Significant gene-based rare variant association results from Sequence Kernel Association Test (SKAT) analyses across ancestry groups and variant classes. Variant classes: Rare variants (MAF < 1%); Potentially Functional, non-intronic variants; Coding, protein-altering coding variants; LoF, predicted gene-disrupting variants; Damaging, missense variants with CADD PHRED ≥ 20; MAF < 0.01% + LoF, loss-of-function variants with MAF < 0.01%; MAF < 0.01% + LoF + Damaging, loss-of-function and damaging missense variants with MAF < 0.01%. Abbreviations: LoF, loss-of-function; MAF, minor allele frequency; N_INFORMATIVE, number of individuals with informative genotype data for the tested region; NumVar, number of variants included in the gene-based test; NumPolyVar, number of polymorphic variants among those included; Q, SKAT test statistic. Abbreviations: AAC, African Admixed; AFR, African; AJ, Ashkenazi Jewish; AMR, American Admixed; CAS, Central Asian; EAS, East Asian; EUR, European; FIN, Finnish; MDE, Middle Eastern; SAS, South Asian; CAH; Complex Admixture.

**Table S3.** Significant gene-based rare variant association results from Sequence Kernel Association Test–Optimal (SKAT-O) analyses across ancestry groups and variant classes. Variant classes: Rare variants (MAF < 1%); Potentially Functional, non-intronic variants; Coding, protein-altering coding variants; LoF, predicted gene-disrupting variants; Damaging, missense variants with CADD PHRED ≥ 20; MAF < 0.01% + LoF, loss-of-function variants with MAF < 0.01%; MAF < 0.01% + LoF + Damaging, loss-of-function and damaging missense variants with MAF < 0.01%. Abbreviations: LoF, loss-of-function; MAF, minor allele frequency; N_INFORMATIVE, number of individuals with informative genotype data for the tested region; NumVar, number of variants included in the gene-based test; NumPolyVar, number of polymorphic variants among those included; Q, SKAT-O test statistic; rho, SKAT-O mixing parameter. Abbreviations: AAC, African Admixed; AFR, African; AJ, Ashkenazi Jewish; AMR, American Admixed; CAS, Central Asian; EAS, East Asian; EUR, European; FIN, Finnish; MDE, Middle Eastern; SAS, South Asian; CAH; Complex Admixture.

## Notes

https://doi.org/10.5281/zenodo.17753486

https://doi.org/10.5281/zenodo.21510483.

### Competing Interest Statement

The authors have declared no competing interest.

### Summary of Updates

missing supplementary tables are now being uploaded.

